# Phylogenetic context using phylogenetic outlines

**DOI:** 10.1101/2021.05.31.446453

**Authors:** Caner Bagci, David Bryant, Banu Cetinkaya, Daniel H. Huson

## Abstract

Microbial studies typically involve the sequencing and assembly of draft genomes for individual microbes or whole microbiomes. Given a draft genome, one first task is to determine its phylogenetic context, that is, to place it relative to the set of related reference genomes. We provide a new interactive graphical tool that addresses this task using Mash sketches to compare against all bacterial and archaeal representative genomes in the GTDB taxonomy, all within the framework of SplitsTree5. The phylogenetic context of the query sequences is then displayed as a phylogenetic outline, a new type of phylogenetic network that is more general that a phylogenetic tree, but significantly less complex than other types of phylogenetic networks. We propose to use such networks, rather than trees, to represent phylogenetic context, because they can express uncertainty in the placement of taxa, whereas a tree must always commit to a specific branching pattern. We illustrate the new method using a number of draft genomes of different assembly quality.

## Introduction

In the study of microbes using sequencing, assembly and contig binning, one important task is to calculate the “phylogenetic context” of a given draft genome, contig, or bin of contigs. This requires that we first determine which known microbes have similar sequences to the query, and then produce a suitable indication of the phylogenetic relationships.

Pairwise distances between genome-scale sequences can be quickly calculated using *k*-mer methods such as Mash [Ondov et al., 2016]. In this type of approach, the *k*-mer content (words of a fixed length *k*) of a sequence is represented by a reduced “sketch” and such sketches are compared using the Jaccard index and derived distance measures that approximate average nucleotide identity (ANI).

The Genome Taxonomy Database (GTDB) [Parks et al., 2020] provides a similarity-based taxonomy for *≈* 195, 000 bacterial and archaeal genomes obtained from the NCBI assembly database [Kitts et al., 2016]. A representative subset of *≈* 32, 000 reference genomes is provided for taxonomic analysis and the GTDB-tk tool kit provides associated analysis tools [Chaumeil et al., 2019].

Here we propose to compute a Mash sketch for each representative reference genome in the GTDB, and to assign a Bloom filter [Bloom, 1970] to each internal node of the taxonomy so as to represent the set of all *k*-mers present in reference genomes below the node [Solomon and Kingsford, 2016, Pierce et al., 2019]. For a given set of query sequences, this will allow one to determine all similar reference genomes quickly enough for use in an interactive program. Mash can then be used to compute a distance matrix on the query and (a subset of) all sufficiently similar references.

Given such a matrix of pairwise distances, one option is to compute a phylogenetic tree to represent the data, using an algorithm such as neighbor-joining [Saitou and Nei, 1987]. Phylogenetic trees are often used to represent such data, because evolution is assumed to be predominantly driven by speciation events. In addition, phylogenetic trees have low complexity, employing only a linear number *O*(*n*) of nodes and edges to represent *n* taxa.

However, in the evolution of microbes, reticulate events, such as horizontal gene transfer and recombination, may play a significant role [Huson et al., 2010]. Also, when using *k*-mer features and distance-based phylogenetic methods, the accuracy of the resulting phylogenetic trees may be poor. Hence, the use of phylogenetic networks, rather than phylogenetic trees, can be more appropriate.

One popular approach to obtaining a phylogenetic network [Huson and Bryant, 2006] is to apply the neighbor-net algorithm [Bryant and Moulton, 2004] on the distances and to represent the output as a splits network [Dress and Huson, 2004], requiring *O*(*n*^4^) nodes and edges, in the worst case.

Here we present a new type of phylogenetic network that we call a *phylogenetic outline*. A phylogenetic outline is also computed from the output of the neighbor-net algorithm and has the mathematical properties of a splits network, thus providing the same information as previously employed networks. A major advantage of phylogenetic outlines is that they are only quadratic in size, containing at most *O*(*n*^2^) nodes and edges. By default, phylogenetic outlines are unrooted, however, we also provide algorithms for both midpoint and out-group rooting.

While our focus here is on using phylogenetic outlines to represent phylogenetic context, please note that phylogenetic outlines can be used to represent the output of the neighbor-net algorithm in all other settings, as well.

The entire procedure described here has been implemented as part of SplitsTree5 (available at: https://software-ab.informatik.uni-tuebingen.de/download/splitstree5). The implementation carries out a Mash comparison of a set of query sequences against a database representing the GTDB, so as to determine the phylogenetic context of the queries, then computes and visualises a phylogenetic outline of the sequences.

Using a single dialog, the user selects the files containing the query sequences, loads a database containing all reference data and then obtains a phylogenetic outline of the queries, interactively in minutes. Unlike other approaches [Ondov et al., 2016, Pierce et al., 2019, Chaumeil et al., 2019], no scripting or running of multiple programs is required.

Conceptually, the calculation of *phylogenetic context* lies between *phylogenetic placement* [Matsen et al., 2010], in which one or more query sequences are placed into a precomputed phylogenetic tree, and *ab initio* phylogenetic tree inference, in which a phylogenetic tree is calculated for all query sequences and a subset of the reference sequences. The GTDB-tk toolkit provides tools for performing phylogenetic placement and *ab initio* tree inference, the latter is performed using a subset of the reference genomes that is delineated by providing an outgroup taxon. In both cases, the result is a phylogenetic tree that can be viewed in a program such as Dendroscope [Huson and Scornavacca, 2012].

To illustrate our method, we apply it to a number of metagenomic draft genomes of different levels of quality, published in [Arumugam et al., 2019]. We also show how this differs from the phylogenetic analyses that one can perform using GTDB-tk.

## Results

Assume that you have sequenced and assembled one or more bacterial genomes, or have calculated a metagenomic binning of contigs. There are a number of command-line pipelines that can be used to determine closely related genomes, ranging from very fast, k-mer based heuristics such as Mash [Ondov et al., 2016], Sourmash [Pierce et al., 2019], or marker-gene based phylogenetic placement methods such as GTDB-tk [Parks et al., 2020], to more thorough, but slower protein-alignment based approaches such as DIAMOND+MEGAN [Buchfink et al., 2015, Huson et al., 2016] or HUMAnN2 [Franzosa et al., 2018]. These methods all require scripting to go from an input file containing one or more sequences of interest to a visualization of the phylogenetic context of the input sequences. Moreover, the visual representation of the context is usually performed using a phylogenetic or taxonomic tree, which presents a definite clustering of taxa with little indication of uncertainty or alternative groupings.

We provide a fast and interactive implementation for exploring phylogenetic context of a set of microbial sequences of interest. The user loads one or more files of query DNA sequences and then requests that all similar reference genomes are determined. Then a threshold is set for the maximum distance of reference genomes, or number of reference genomes, to be considered. These are downloaded and a Mash comparison of the query sequences and all similar reference genomes is performed, the neighbor-net method is run and the result is presented as a phylogenetic outline. (The user can also choose to use a tree-building method such as neighbor-joining [Saitou and Nei, 1987]).

To illustrate our method, we applied it to a number of draft genomes reported in [Arumugam et al., 2019]. These draft genomes, or metagenomic assembly bins, contain assembled contigs of long-read microbiome sequences obtained from a bio-reactor enriched for polyphosphate accumulation. The paper reports a taxonomic assignment for each bin that is based on an analysis of the contained protein-coding genes and confirmed using 16S rRNA sequences, when present. For each of the 14 reported bins, we calculated a phylogenetic outline to display the phylogenetic context of the closest reference genomes below a certain distance. Three are displayed in Figure 1 (one “high-quality”, one “medium-quality” and one “low-quality” draft genome, respectively, as defined in [Bowers et al., 2017]) and the other 11 are show in the Supplement.

**Figure 1.**
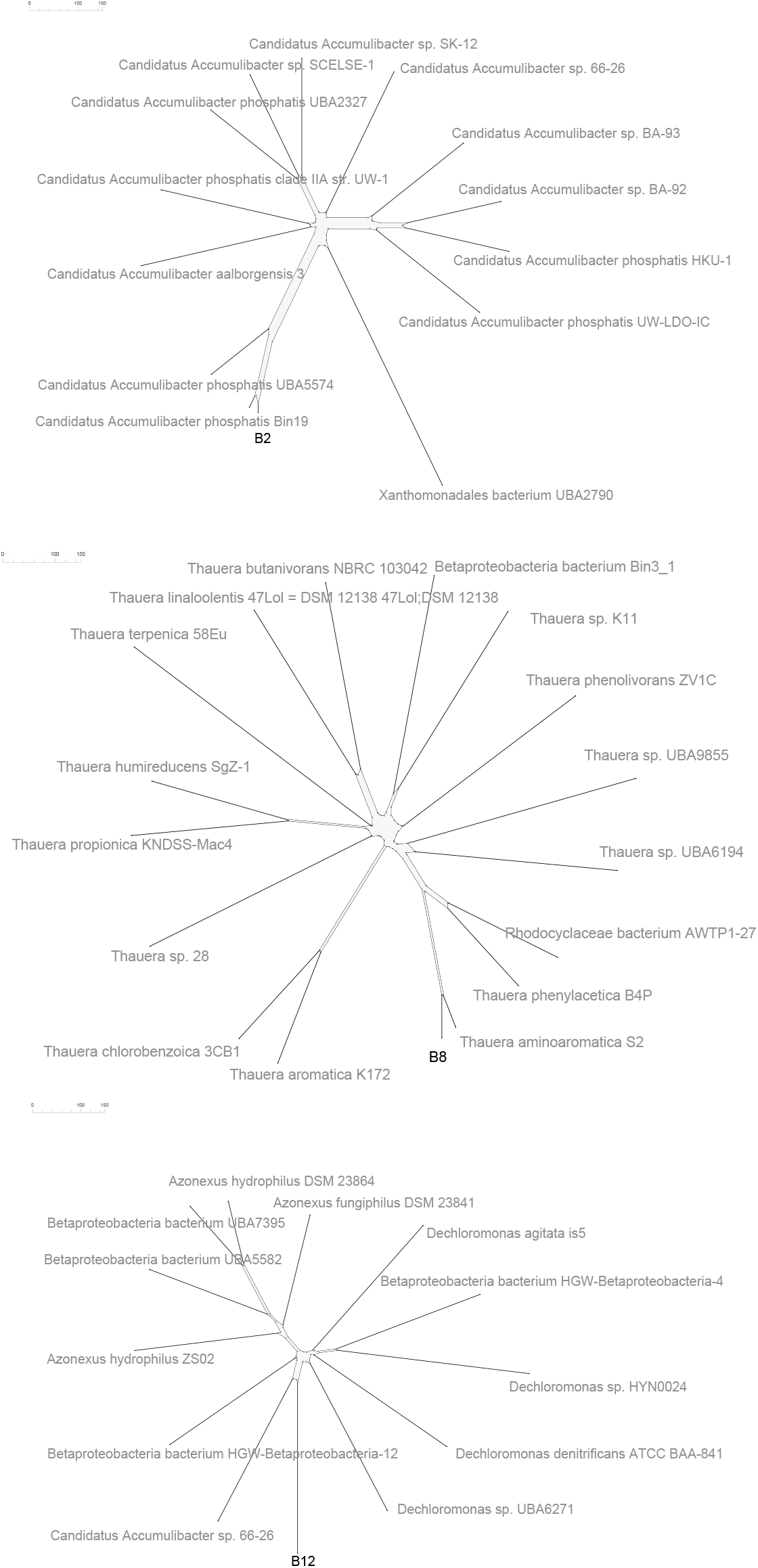
Phylogenetic context. The phylogenetic outlines computed by SplitsTree5 for three metagenomic draft genomes presented in [Arumugam et al., 2019], (B2, B8, and B12).

Generally speaking, in all three cases, the phylogenetic context is compatible with the reported taxonomic identity.

In the case of draft genome B2, all (but one) reference genomes displayed in the phylogenetic context are members of the genus *Candidatus Accumulibacter*. This is in agreement with the classification presented in [Arumugam et al., 2019], which assigned B2 to the species *Candidatus Accumulibacter* sp. SK-02. The closest species in the phylogenetic context analysis is *Candidatus Accumulibacter phosphatis* Bin19, Mash distance 0.01, a genome that was not available to Arumugam et al. [2019]. The second closest, *Candidatus Accumulibacter phosphatis* UBA5574, Mash distance 0.07, is not represented by protein sequences in NCBI and was thus was not part of the database used by Arumugam et al. [2019].

Unexpectedly, the species *Xanthomonadales bacterium* UBA2790, which comes from a different taxonomic class, also appears in the phylogenetic context of B2, with a Mash distance of 0.2. Note that this metagenome assembled genome (MAG) comes from a sample of granular sludge that also gave rise to two *Candidatus Accumulibacter* reference genomes [Parks et al., 2017] and we suspect that it might be contaminated with *Candidatus Accumulibacter* contigs or sequence.

In the case of draft genome B8, all references genomes displayed in the phylogenetic context are *Thauera* species, except one unclassified Betaproteobacteria bacterium, which is, however, a member of the genus *Thauera*, and this supports the assignment to the genus *Thauera*. The closest reference genome *Thauera aminoaromatica* S2 has a Mash distance of 0.2.

Finally, in [Arumugam et al., 2019], the draft genome B12 was classified as a member of the Betaproteobacteria class, suggesting that there did not exist a closely related reference at the time of the publication of the dataset. In the phylogenetic context computed by SplitsTree, B12 is placed closest to *Candidatus Accumulibacter* sp. 66-26 with a Mash distance of 0.14. In addition, some species of the genera *Azonexus* and *Dechloromonas* can be found that are similar to B12, with Mash distances below 0.18.

These two genera belong to the family of Azonexaceae in the NCBI taxonomy, while *Candidatus Accumulibacter* does not have a defined family or order. All three genera belong to the family Rhodocyclaceae in the GTDB taxonomy. Although B12 does not have many closely related reference genomes, the phylogenetic outline produced by SplitsTree5 suggests that B12 belongs to the Rhodocyclaceae family, which is more specific that the assignment suggested by Arumugam et al. [2019].

In each of the three examples, it took between one and three minutes to determine all reference genomes whose sketches have a distance of at most 0.3 to the sketch of the draft genome, and then to compute and display the phylogenetic outlines for the 10 most similar references. Computations were carried out on a laptop with 8 cores (at 2.4 GHz) and 32GB of memory. Reference genomes are downloaded (and cached) on demand, which takes additional time. The distance thresholds used to select the closest reference genomes for each bin were chosen interactively and are reported in the Supplementary file.

To illustrate the improved practical performance of the outline algorithm on a larger dataset, we computed the phylogenetic context for bin B12 using the 1,000 closest reference genomes. Running the neighbor-net algorithm on this data takes 90 seconds and results in 4,516 splits. The equal-angle algorithm [Dress and Huson, 2004] produces a splits network with 108,640 nodes and 212,762 edges, and requires about seven minutes to compute and show the network. In contrast, our new outline algorithm produces a splits network with 8,028 nodes and 8,028 edges and requires only two seconds for this (not shown here).

For the purpose of comparison, we applied GTDB-Tk [Chaumeil et al., 2019] in phylogenetic placement mode (`classify wf` workflow) to the draft genomes B2, B8 and B12. GTDB-Tk uses GTDB accessions to label reference genomes, whereas SplitsTree uses the strain names associated with the assemblies in the assembly reports of NCBI. In Figure 2, we assign the same color to corresponding GTDB accessions and strain names.

**Figure 2.**
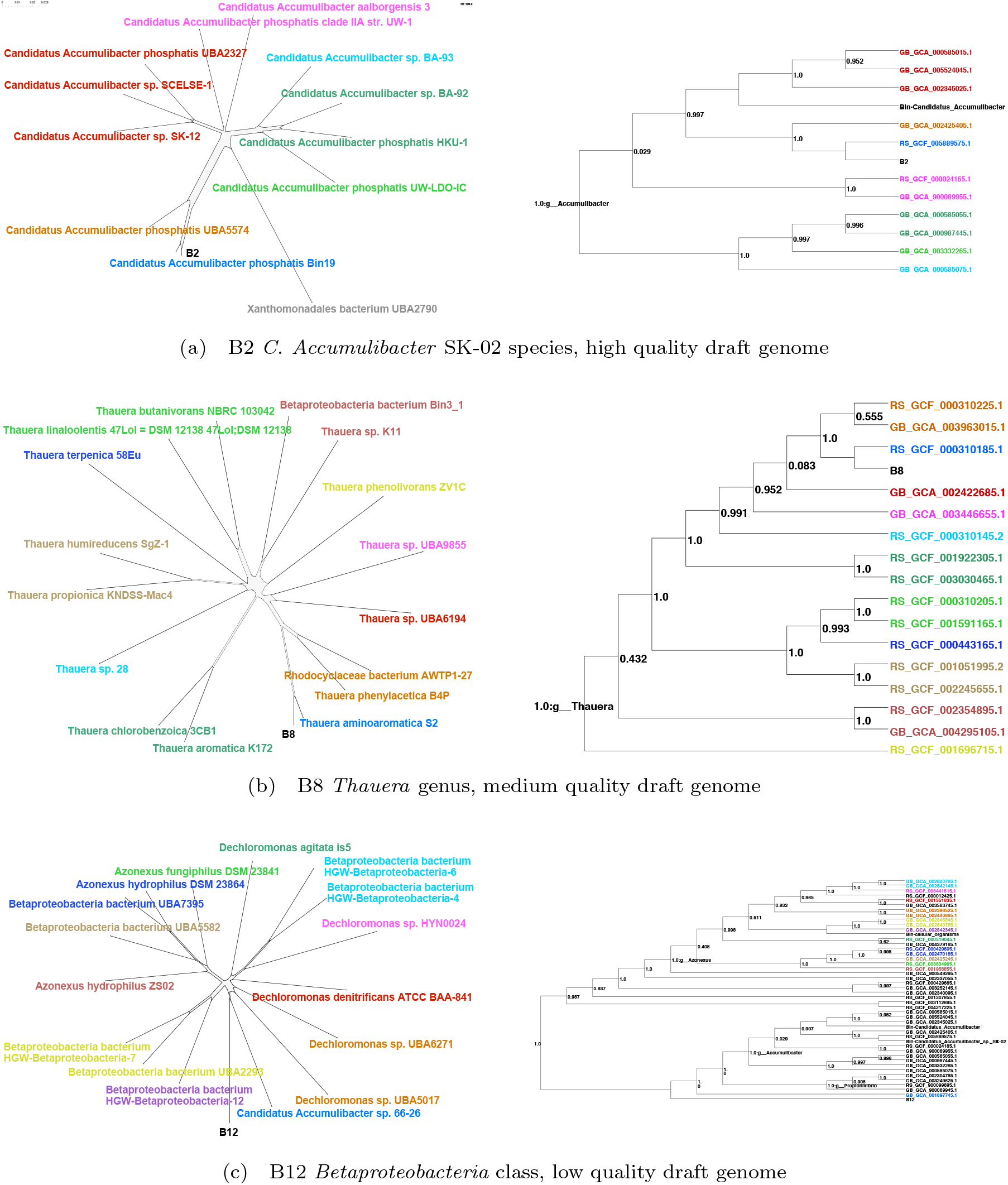
Phylogenetic context. For three metagenomic draft genomes B2, B8, and B13, we report the taxonomic assignment and genome quality [Arumugam et al., 2019], and display both the phylogenetic outline computed by SplitsTree5 and a tree representing the phylogenetic placement computed using GTDB-Tk.

In the case of draft genome B2, the tree computed by GTDB-Tk agrees very well with the phylogenetic context computed using SplitsTree, placing B2 next to *Candidatus Accumulibacter phosphatis* Bin19, and to other reference genomes shown in the phylogenetic outline, see Figure 2a. The distances computed by SplitsTree5 was also similar to those reported by GTDB-Tk.

In the case of draft genome B8, the phylogenetic context included all members of the genus *Thauera* from GTDB-Tk, and placed the query bin next to *Thauera aminoaromatica* S2. The tree produced by GTDB-Tk contains the same references and has a similar topology. See Figure 2b. This suggests that, if distances between genomes are small, then a Mash-based analysis, as in SplitsTree5, may perform very similar to a marker-gene and ANI based analysis, as in GTDB-Tk.

Finally, in the case of B12, this draft genome is further away from any reference genome than the two draft genomes just discussed. See Figure 2c. GTDB-Tk places B12 outside of the boundaries of any genera, but closer to the genus *Accumulibacter*, and closest to the species *Candidatus Accumulibacter* sp. 66-26.

SplitsTree5 also places B12 closest to the species *Candidatus Accumulibacter* sp. 66-26; however, the rest of the references shown in the phylogenetic context are from the genus *Azonexus* instead of *Accumulibacter*. Although the distances within the genus are in agreement with those computed by GTDB, here we see a difference in the phylogeny outside genus boundaries.

We also determined the phylogenetic context and GTDB-Tk placement for all other 11 MAGs reported in [Arumugam et al., 2019] and report these in the Supplementary File. For those draft genomes for which very similar reference genomes can be found, the phylogenetic context computed by SplitsTree is similar to the phylogenetic placement computed by GTDB-Tk. In the other cases, either the phylogenetic context contains only very few references, or it contains a wide range of different references and disagrees with the phylogenetic placement computed by GTDB-Tk (see B4 and B6 in the Supplementary File). These disagreements persist even if one uses a more accurate calculation of average nucleotide identity (not shown here), indicating that they are due to a fundamental difference between ANI analysis and marker-gene analysis.

## Materials and Methods

### Preprocessing the reference database

We downloaded the GTDB taxonomy [Parks et al., 2020] in July 2020. The taxonomy has 240,103 nodes, of which 194,600 are leaves. GTDB identifies 31,910 genomes representative genomes. These are available from the GTDB download page https://data.gtdb.ecogenomic.org/releases/latest/genomic_files_reps/. Links to the other (non-representative) genomes are contained in the GenBank or RefSeq [Pruitt et al., 2009] assembly summary reports on the NCBI genomes FTP site ftp://ftp.ncbi.nlm.nih.gov/genomes/ASSEMBLY_REPORTS/.

In a processing step, we computed a Mash sketch [Ondov et al., 2016] for each of the 31,910 representative genomes, using a word size of *k* = 21 and sketch size of *s* = 10, 000. Multi-part genome sequences were concatenated. For each internal node of the GTDB taxonomy, we computed a Bloom filter [Bloom, 1970] representing all *k*-mers contained in all sketches associated with genomes below the node, using a false positive probability of 0.0001. For these calculations, we used our own implementations of the Mash algorithm, `mash-sketches`, and Bloom filters, `bfilter-tool`, which we provide as a part of our SplitsTree5 package.

All taxa, Mash sketches, Bloom filters and genome URL’s were loaded into an SQLITE [Hipp, 2020] database file. In addition, the file contains an explicit representation of the GTDB taxonomy using a node-to-parent mapping. The database schema is shown in Figure 3. The database file is 12.4Gb in size and does not contain the actual genome sequences; these are downloaded (and cached) by our implementation on demand.

**Figure 3.**
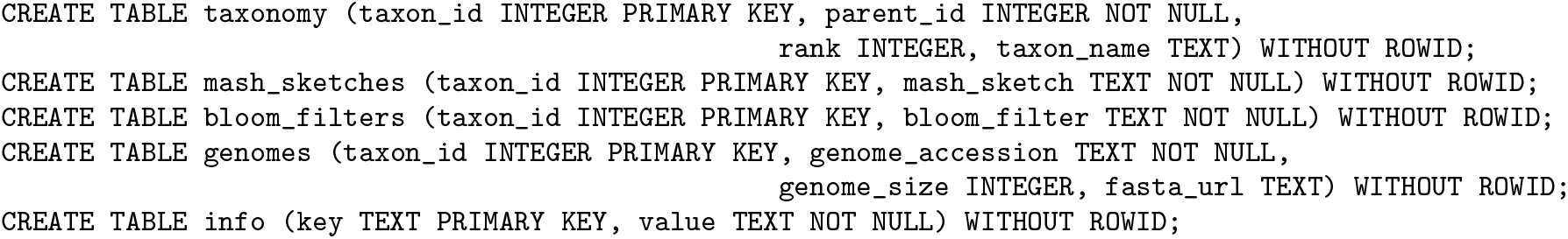
Database schema. The `taxa` table contains the taxon id, name and the id of the parent node in the taxonomy. The `mask sketches` and `bloom filters` tables contain all Mash sketches and Bloom filters. The `genomes` table contains the genome accession, genome size and an URL of a FastA file containing the genome sequence. Finally, the `info` table contains general information such as version and size of the database.

SplitsTree5 and the current database file `gtdb-rep-k21-s10000-May2021.db` can can be downloaded here: https://software-ab.informatik.uni-tuebingen.de/download/splitstree5.

### The outline algorithm

For a given distance matrix *D* on a set of *n* taxa *𝒳*, the neighbor-net algorithm [Bryant and Moulton, 2004] computes a set of weighted splits Σ of *𝒳*, that is, a set of bipartitions of the form *S* = *A* | *B*, where *A* ≠ ∅, *B* ≠ ∅, *A* ∩ *B* = ∅ and *A* ∪ *B* = *𝒳*. The set of splits computed by neighbor-net has quadratic size *O*(*n*^2^). The set of splits is “circular”, which implies that they can be represented by an “outer-labeled planar” splits network [Dress and Huson, 2004], see Figure 4(b), using *O*(*n*^4^) nodes and edges, in the worst case.

**Figure 4.**
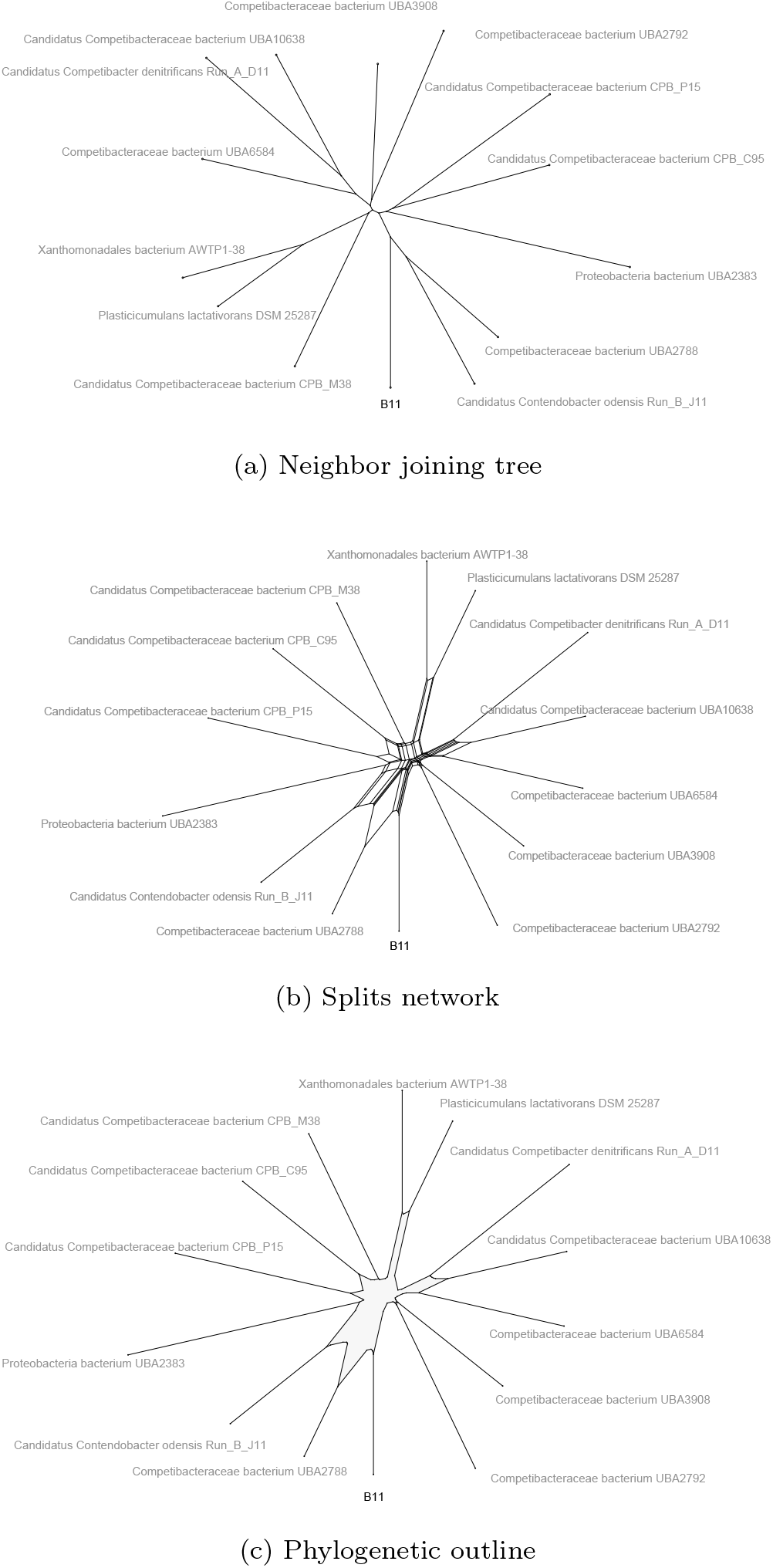
Tree and networks. For a low quality draft genome “B11” from [Arumugam et al., 2019], we display its calculated phylogenetic context, using (a) a neighbor-joining tree with 26 nodes and 25 edges, (b) a splits network with 120 nodes and 197 edges, and (c) a phylogenetic outline with 68 nodes and 68 edges, respectively.

Here we describe the computation of a phylogenetic outline that requires only *O*(*n*^2^) nodes and edges, see Figure 4(c). In a phylogenetic outline, each split *S* = *A* | *B* is represented by a single edge, or two parallel edges, that separate all taxa in *A* from all taxa in *B*, and thus a phylogenetic outline fulfills the definition of a splits network [Dress and Huson, 2004].

Consider a set Σ of *m* splits on *𝒳*, each split *S* with a positive weight *ω*(*S*). Assume, without loss of generality, that Σ contains all trivial splits on *𝒳*, that is, all splits that separate exactly one taxon from all others. We will assume that the splits are circular, that is, that there exists an ordering *x*_1_, *x*_2_, …, *x*_*n*_ of the taxon set *𝒳* such that each split *S* ∈ Σ can be written as *S* = *{x*_*i*_, …, *x*_*j*_*}* | *𝒳* − *{x*_*i*_, …, *x*_*j*_}, with 1 < *i j n*, in other words, as an interval of elements of *𝒳*, which does not contain the first taxon, vs all others. This condition is always satisfied by the output of neighbor-net [Bryant and Moulton, 2002].

To illustrate this, consider the set of splits 𝒮 = {*S*_1_, …, *S*_5_, *S*_*a*_, *S*_*b*_, *S*_*c*_} on *𝒳* = {*x*_1_, …, .*x*_5_}, where *S*_*a*_ = {*x*_2_, *x*_3_} | {*x*_1_, *x*_4_, *x*_5_}, *S*_*b*_ = {*x*_3_, *x*_4_, *x*_5_} | {*x*_1_, *x*_2_} and *S*_*c*_ = *{x*_3_, *x*_4_} | *{x*_1_, *x*_2_, *x*_5_}. Moreover, for *i* = 1, … 5, let *S*_*i*_ be the trivial split separating *x*_*i*_from all other taxa. This set of splits is circular, as illustrated in Figure 5a.

**Figure 5.**
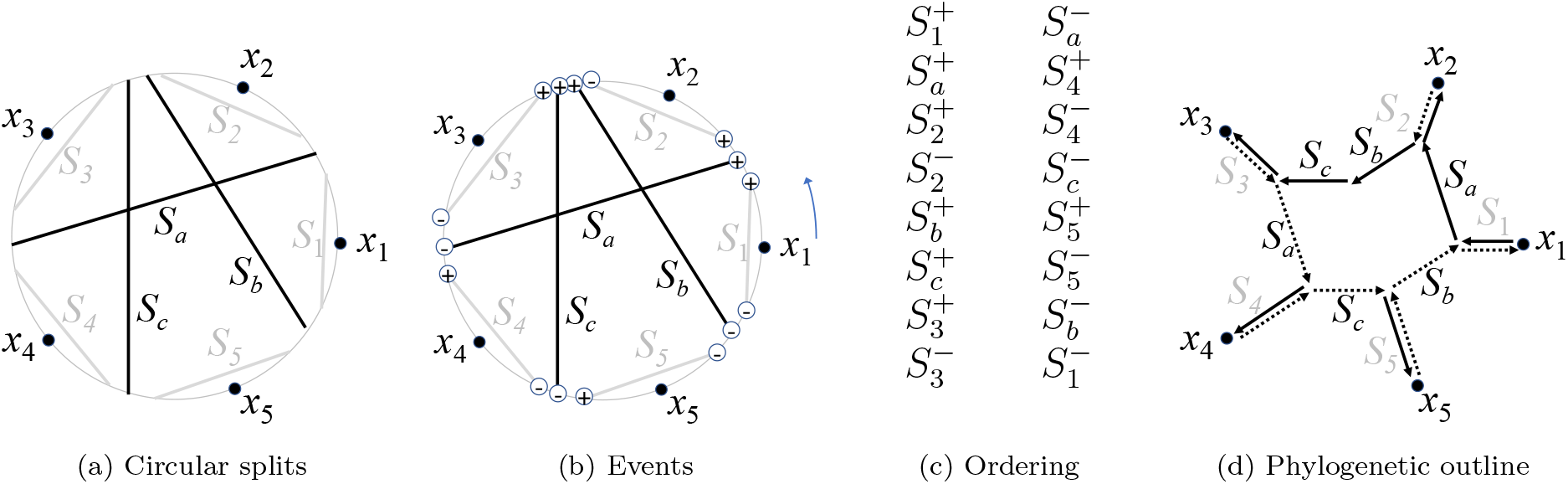
Circular splits and outline. (a) A set of splits {*S*_1_, …, *S*_5_, *S*_*a*_, *S*_*b*_, *S*_*c*_} that is circular, that is, for which the taxa can be placed around a circle such all splits correspond to chords of the circle. (b) Travelling around the circle in positive orientation, each split *S* is encountered twice, first where the interval that does not contain taxon *x*_1_ starts (outbound event *S*^+^ marked ⊕) and then again where that interval ends (inbound event *S*^−^ marked 8). ⦵) (c) The ordered events, listed in two columns. (d) Starting at *x*_1_, in the order of the events, when encountering an outbound event *S*^+^ move perpendicularly to the chord for split *S* by a distance of *ω*(*S*), as indicated by solid arrows. When encountering an inbound event *S*^−^, move in the opposite direction by the same distance, as indicated by dotted arrows.

Circularity implies that, for each split *S* ∈ Σ, the split part not containing *x*_1_ is an interval of the form *I*(*S*) = *{x*_*i*_, …, *x*_*j*_} with 1 < *i* ⩽ *j* ⩽ *n*. We will use *i*(*S*) and *j*(*S*) to refer to the two interval bounds.

Our new *outline algorithm* for computing a phylogenetic outline proceeds in three steps. In summary, first, we define two “events” per split. Second, we sort all events. Third we process all events in sorted order, constructing either 0 or 1 new nodes and/or edges, per event.

For each split *S* we define two events, an *outbound event S*^+^, crossing over to the other side of *S* from the side that contains *x*_1_, and an *inbound event S*^−^, returning back to the side of *S* that contains *x*_1_. We will sort these events and then use them to construct the phylogenetic outline.

We define a total ordering on all events as follows (see Figure 5b-c):

- For two outbound events *S*^+^ and *T* ^+^, set *S*^+^ < *T* ^+^, if either *i*(*S*) < *i*(*T*) or both *i*(*S*) = *i*(*T*) and *j*(*S*) > *j*(*T*).
- For two inbound events *S*^−^ and *T* ^−^, set *S*^−^ < *T* ^−^, if either *j*(*S*) < *j*(*T*) or both *j*(*S*) = *j*(*T*) and *i*(*S*) > *i*(*T*).
- For an outbound event *S*^+^ and an inbound event *T* ^−^, set *S*^+^ < *T* ^−^, if *i*(*S*) < *j*(*T*) + 1, and set *S*^+^ > *T* ^−^, otherwise.

The ordering of all *O*(*n*^2^) events can be computed in *O*(*n*^2^) steps: Use radix sort to first sort all outbound events *S*^+^ in decreasing order of *j*(*S*), and then in increasing order of *i*(*S*). Similarly, use radix sort to first sort all inbound events *S*^−^ in decreasing order of *i*(*S*) and then in increasing order of *j*(*S*). Finally merge the two lists of events observing the relative ordering of outbound and inbound events.

We now describe how to create the nodes and edges of the outline (see Figure 5d).

We will use *p* to denote the current location, initially set to (0, 0). Place taxon *x*_1_ on a new node *v*_1_ at location *π*(*v*_1_) = *p*. We will use Σ(*v*) to denote the set of splits that separates a node *v* from the node *v*_1_.

For each split *S*, we define an angle

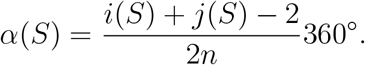

In the example in Figure 5b, this is perpendicular to the chord associated with *S*.

We process all events as described in the following two paragraphs. Let *v* be the current node, initially set to *v*_1_.

To process an outbound event for a split *S*, move the current location *p* in the direction of *α*(*S*) by a distance of *ω*(*S*), the given positive weight of *S*. Create a new node *w* and connect *v* to *w* by a new edge. Set Σ(*w*) = Σ(*v*) *∪* {*S*}. Update the current node, setting *v* = *w*.

To process an inbound event for a split *S*, move the current location *p* in the opposite direction of *α*(*S*) by a distance of *ω*(*S*). Consider the set of splits Σ = Σ(*v*) − *{S}*. We set *w* = *u*, if there exists a node *u* with Σ(*u*) = Σ′. Else, we create a new node *w*, and set *π*(*w*) = *p* and Σ(*w*) = Σ′. We connect *v* and *w* by an edge, if they are not already connected by an edge. Update the current node, setting *v* = *w*.

The number *m* of circular splits on *n* taxa is bounded by *O*(*n*^2^). As discussed above, there will be at most 2*m* nodes and 2*m* edges in the network, and therefore the size is bounded by *O*(*n*^2^). The events are sorted using radix sort, in time linear in the number of events, and thus in *O*(*n*^2^) time. The construction of nodes and edges also requires only *O*(*n*^2^) steps. Hence, the outline algorithm requires at most *O*(*n*^2^) in total. The network size and time requirement compare favorably to the *O*(*n*^4^) network size and time worst-case requirements of the equal angle algorithm [Dress and Huson, 2004], which is currently used to visualize the output of the neighbor-net algorithm in SplitsTree4 [Huson and Bryant, 2006].

We now discuss how to compute a rooted phylogenetic outline. For midpoint rooting, we proceed as follows. We first determine two taxa, *a* and *b*, that maximize the split distance *d*_Σ_(*a, b*) = _*S*∈Σ(*a,b*)_ *ω*(*S*), where the sum is taken over the set Σ(*a, b*) of all splits *S* that separate *a* and *b*. The set Σ(*a, b*) is then sorted by increasing cardinality of the split part *S*(*a*) containing *a*, and then by increasing size of the intersection of *S*(*a*) with the interval of all taxa that lie between *a* and *b* in the cycle. The root is then positioned in the first split for which the accumulated sum of weights is at least half of *d*_Σ_(*a, b*). For rooting by out-group, the root is placed in the middle of a split that separates the out-group from the rest of the taxa and is minimal with respect to that property.

### Graphical user interface

We have implemented the approach described here in our program SplitsTree5. To compute a phylogenetic outline displaying the phylogenetic context for one or more prokaryotic sequences, select the `File` → `Analyze Genomes…` menu item. This will open a dialog with three tabs. The first tab is used to select the input file(s) and output file, and to determine whether all sequences in a given file are to be concatenated, or are to be treated separately (see Figure 6a). In addition, one can set a minimum sequence length (here set to 100,000 bp). The second tab is used to edit the names for the sequences (see Figure 6b). The third tab is used to perform a Mash-based search in the GTDB database and to select which reference genomes should be included in the phylogenetic outline, based on their distances to the input sequences (see Figure 6c).

**Figure 6.**
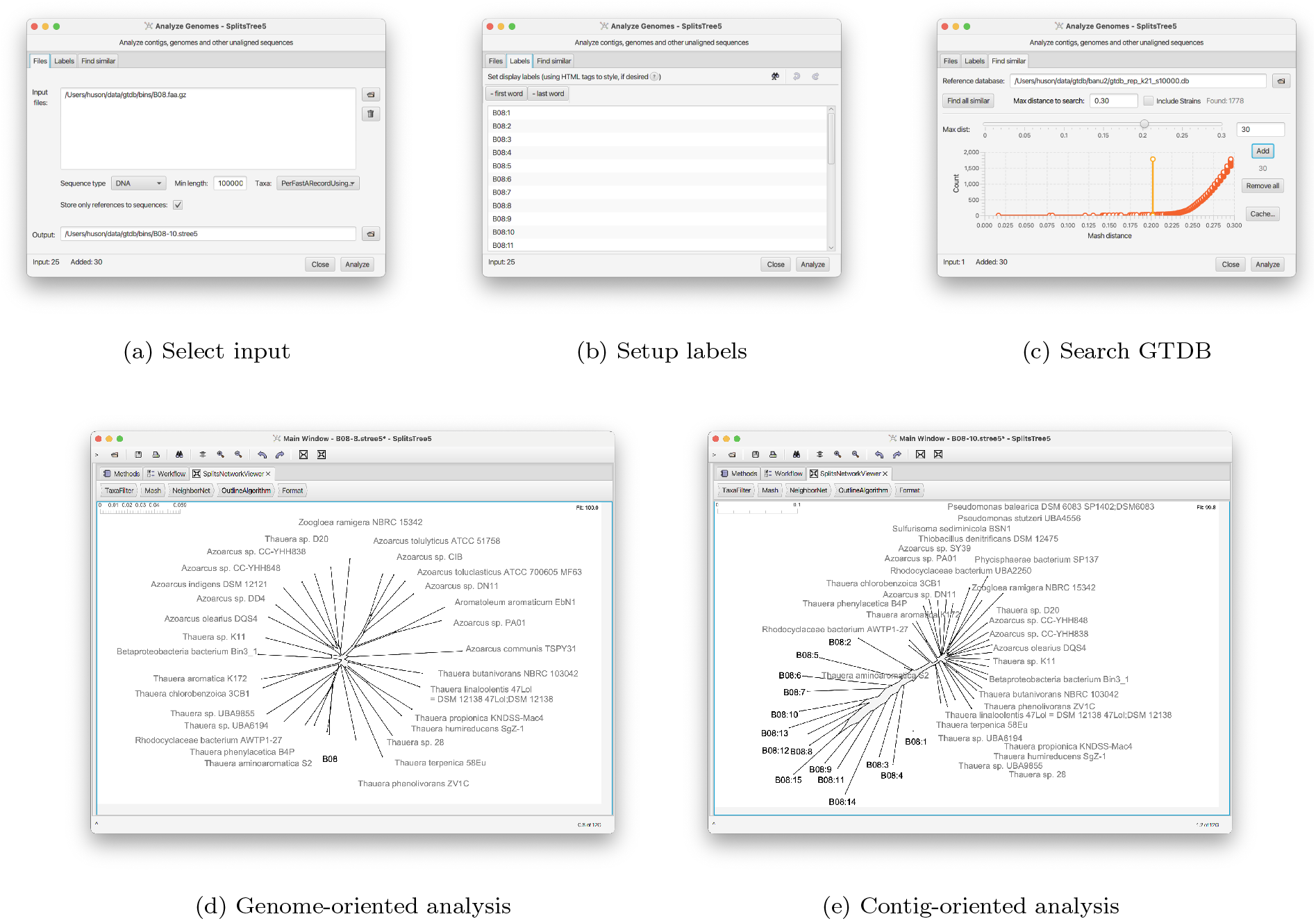
Phylogenetic context analysis. (a) The user selects the input file(s) and decides whether to analyze on a “per file” (complete genome) or “per FastA record” (individual contigs) basis. (b) The labels are set for the input sequences. (c) A search against the GTDB database is initiated and a threshold for the maximum distance is set. Once completed, a phylogenetic outline is drawn. (d) In a genome-oriented analysis, the phylogenetic outline shows the context of the concatenated input sequences. (e) Alternatively, in a contig-oriented analysis, the different sequences in the input file are represented individually in the phylogenetic outline.

The example presented is a medium quality draft genome consisting of 25 contigs assembled from long read sequences, designated bin B8 in [Arumugam et al., 2019] with taxonomic assignment to the genus *Thauera*. In Figure 6d we use a phylogenetic outline to show the phylogenetic context of the draft genome, involving the 30 closest reference genomes.

In Figure 6e we show the phylogenetic context for the 15 (of 25) input contigs whose length achieves the set threshold of 100,000 bp. The contigs are numbered by decreasing length, ranging from B08:1 with length 770,679 bp to B08:15 with length 109,403 bp, respectively. While the outline indicates that the longest contig B08:1 is very similar to the shown reference genomes, the similarity between contigs and references decreases with decreasing contig length.

In addition, we provide a Python implementation of neighbor-net and phylogenetic outlines here: https://github.com/husonlab/SplitsPy.

### Running GTDB-Tk

The frame-shift corrected bins from [Arumugam et al., 2019] were classified using the phylogenetic placement mode of GTDB-Tk [Chaumeil et al., 2019], using the GTDB database R95 version [Parks et al., 2020]. We ran the `classify wf` workflow with the default settings, using 32 cores both for the main pipeline and for pplacer. GTDB-Tk completed the phylogenetic placement of all bins in 26 minutes. In order to visualize the resulting phylogenetic placements, we opened the Newick-formatted `gtdbtk.bac120.classify.tree` output file in Dendroscope [Huson and Scornavacca, 2012] and manually extracted the relevant subtrees for the bins shown in Figure 2 and the Supplementary File.

## Discussion

Here we bring together a number of different ideas, using the GTDB database to represent the taxonomy of bacterial and archaeal genomes; Mash sketches and Bloom filters for fast sequence comparison; and the neighbor-net method and our new concept of phylogenetic outlines for visualization. We thus provide a fast heuristic for establishing the phylogenetic context for one or more prokaryotic genomes or DNA sequences. We demonstrated that our approach can be applied to usefully determine and visualize the phylogenetic context of bacterial draft genomes at different levels of assembly quality.

We believe that the use of a phylogenetic outline, rather than a phylogenetic tree, to represent phylogenetic context is more suitable because outlines can express vagueness in the placement of taxa with respect to each, whereas trees suggest a specific branching pattern. For example, in Figure 4a we show the unrooted, resolved phylogenetic tree computed using the neighbor-joining algorithm [Saitou and Nei, 1987]. Both the splits network (Figure 4b) and the phylogenetic outline (Figure 4c) place *Competibacteraceae bacterium UBA2788* halfway between *Candidatus Contendobacter odensis Run B J11* and the draft genome B11. This ambiguity of placement is not evident in the tree representation.

The mash-based calculation of phylogenetic outlines presented here is, on the one hand, a form of *ab initio* phylogenetic analysis, in which we infer evolutionary relationships from data. On the other hand, the aim is to visualize the phylogenetic context of sequences, which is similar to the goal of phylogenetic placement. We provide a graphical user interface that allows the user to interactively compute and explore the context of sequences. While GTDB-tk provides methods for both phylogenetic placement and ab initio phylogenetic analysis, these calculations are performed using a script and the output is presented as text files. While the resulting trees can viewed using third party tools, the leaves of the tree are labeled by GTDB accessions, whereas our approach provides the option of labeling leaves by the associated NCBI names.

Using a phylogenetic outline to represent phylogenetic context does not replace careful alignment and sophisticated phylogenetic analysis when the goal is to understand the evolutionary history of a set of taxa in detail. Nevertheless, we believe that our approach will prove to be a useful addition to the biologists’ computational toolbox.

## Contributions

DB and DHH conceptualized the project. DB and DHH developed the outline algorithm. DHH designed and implemented the software. CB and BC designed and populated the database. CB performed all data analysis and comparisons. DHH and CB wrote the original draft of the manuscript and all authors edited the manuscript.

## Acknowledgements

DHH acknowledges Catalyst: Leaders funding provided by the New Zealand Ministry of Business, Innovation and Employment and administered by the Royal Society Te Apārangi. The authors acknowledge infrastructural support by the cluster of Excellence EXC2124 Controlling Microbes to Fight Infection (CMFI), project ID 390838134

## Notes

### Competing Interest Statement

The authors have declared no competing interest.

## References

K. Arumugam, C. Bagci, I. Bessarab, S. Beier, B. Buchfink, A. Gorska, G. Qiu, D.H. Huson, and R.B.H. Williams. Annotated bacterial chromosomes from frame-shift-corrected long read metagenomic data. Microbiome, 7(61), 2019.

B.H. Bloom. Space/time trade-offs in hash coding with allowable errors. Commun. ACM, 13(7):422–426, July 1970.

Robert M Bowers, Nikos C Kyrpides, Ramunas Stepanauskas, Miranda Harmon-Smith, Devin Doud, T B K Reddy, Frederik Schulz, Jessica Jarett, Adam R Rivers, Emiley A Eloe-Fadrosh, Susannah G Tringe, Natalia N Ivanova, Alex Copeland, Alicia Clum, Eric D Becraft, Rex R Malmstrom, Bruce Birren, Mircea Podar, Peer Bork, George M Weinstock, George M Garrity, Jeremy A Dodsworth, Shibu Yooseph, Granger Sutton, Frank O Glöckner, Jack A Gilbert, William C Nelson, Steven J Hallam, Sean P Jungbluth, Thijs J G Ettema, Scott Tighe, Konstantinos T Konstantinidis, Wen-Tso Liu, Brett J Baker, Thomas Rattei, Jonathan A Eisen, Brian Hedlund, Katherine D McMahon, Noah Fierer, Rob Knight, Rob Finn, Guy Cochrane, Ilene Karsch-Mizrachi, Gene W Tyson, Christian Rinke, Lynn Schriml, Philip Hugenholtz, Pelin Yilmaz, Folker Meyer, Alla Lapidus, Donovan H Parks, A. Murat Eren, Jillian F Banfield, Tanja Woyke, and The Genome Standards Consortium. Minimum information about a single amplified genome (misag) and a metagenome-assembled genome (mimag) of bacteria and archaea. Nature Biotechnology, 35(8):725–731, 2017.

D. Bryant and V. Moulton. NeighborNet: An agglomerative method for the construction of planar phylogenetic networks. In R. Guigó and D. Gusfield, editors, Algorithms in Bioinformatics, WABI 2002, volume LNCS 2452, pages 375–391, 2002.

D. Bryant and V. Moulton. Neighbor-net: An agglomerative method for the construction of phylogenetic networks. Molecular Biology and Evolution, 21(2):255–265, 2004.

B. Buchfink, C. Xie, and D.H. Huson. Fast and sensitive protein alignment using DIAMOND. Nature Methods, 12:59–60, 2015.

P.-A. Chaumeil, A.J. Mussig, P. Hugenholtz, and D.H. Parks. GTDB-Tk: a toolkit to classify genomes with the genome taxonomy database. Bioinformatics, 36(6):1925–1927, 11 2019.

A.W.M. Dress and D.H. Huson. Constructing splits graphs. IEEE/ACM Transactions in Computational Biology and Bioinformatics, 1(3):109–115, 2004.

E.A. Franzosa, L.J. McIver, G. Rahnavard, L.R. Thompson, M. Schirmer, G. Weingart, K. Schwarzberg Lipson, R. Knight, J.G. Caporaso, N. Segata, and C. Huttenhower. Species-level functional profiling of metagenomes and metatranscriptomes. Nature methods, 15(11):962–968, 11 2018. doi: 10.1038/s41592-018-0176-y.

Richard D Hipp. SQLite, 2020. URL https://www.sqlite.org/index.html.

D. H. Huson and C. Scornavacca. Dendroscope 3 - a program for computing and drawing rooted phylogenetic trees and networks. Systematic Biology, 61(6):1061–7, 2012.

D.H. Huson and D. Bryant. Application of phylogenetic networks in evolutionary studies. Molecular Biology and Evolution, 23:254–267, 2006.

D.H. Huson, R. Rupp, and C. Scornavacca. Phylogenetic Networks. Cambridge University Press, 2010.

D.H. Huson, Sina Beier, Isabell Flade, Anna Górska, Mohamed El-Hadidi, Suparna Mitra, Hans-Joachim Ruscheweyh, and Rewati Tappu. MEGAN Community Edition - interactive exploration and analysis of large-scale microbiome sequencing data. PLoS Comput Biol, 12(6):e1004957, 2016. doi:10.1371/journal.pcbi.1004957.

P.A. Kitts, D.M. Church, F. Thibaud-Nissen, J. Choi, V. Hem, V. Sapojnikov, R.G. Smith, T. Tatusova, C. Xiang, A. Zherikov, M. DiCuccio, T.D. Murphy, K.D. Pruitt, and A. Kimchi. Assembly: a resource for assembled genomes at ncbi. Nucleic Acids Res, 44 (D1):D73–80, Jan 2016.

F.A. Matsen, R.B. Kodner, and E.V. Armbrust. pplacer: linear time maximum-likelihood and bayesian phylogenetic placement of sequences onto a fixed reference tree. BMC bioinformatics, 11(1):538+, October 2010.

B.D. Ondov, T.J. Treangen, P. Melsted, A.B. Mallonee, N.H. Bergman, S. Koren, and A.M. Phillippy. Mash: fast genome and metagenome distance estimation using MinHash. Genome Biology, 17(1):132, 2016.

D.H. Parks, M. Chuvochina, P.-A. Chaumeil, C. Rinke, A.J. Mussig, and P. Hugenholtz. A complete domain-to-species taxonomy for bacteria and archaea. Nature Biotechnology, 2020. doi: 10.1038/s41587-020-0501-8.

Donovan H Parks, Christian Rinke, Maria Chuvochina, Pierre-Alain Chaumeil, Ben J Woodcroft, Paul N Evans, Philip Hugenholtz, and Gene W Tyson. Recovery of nearly 8,000 metagenome-assembled genomes substantially expands the tree of life. Nature microbiology, 2(11):1533–1542, November 2017. ISSN 2058-5276. doi: 10.1038/s41564-017-0012-7. URL http://dx.doi.org/10.1038/s41564-017-0012-7.

N.T. Pierce, L. Irber, T. Reiter, P. Brooks, and C.T. Brown. Large-scale sequence comparisons with sourmash. F1000Research, 8, 2019.

K.D. Pruitt, T. Tatusova, W. Klimke, and D.R. Maglott. NCBI reference sequences: current status, policy and new initiatives. Nucleic Acids Res., pages D32–36, 2009.

N. Saitou and M. Nei. The Neighbor-Joining method: a new method for reconstructing phylogenetic trees. Molecular Biology and Evolution, 4:406–425, 1987.

B. Solomon and C. Kingsford. Fast search of thousands of short-read sequencing experiments. Nat Biotechnol, 34(3):300–302, Mar 2016.

