## Supplementary File for "Phylogenetic context using phylogenetic outlines"

### Supplementary Material

Table S1: Number of genomes included in the phylogenetic context of each metagenomic bins according to the max distance parameter that was chosen

| **Bin** | **# genomes** | **Max distance** |
|---|---|---|
| B1 | 6 | 0.3 |
| B2 | 13 | 0.25 |
| B3 | 2 | 0.3 |
| B4 | 7 | 0.275 |
| B5 | 4 | 0.3 |
| B6 | 16 | 0.28 |
| B7 | 2 | 0.3 |
| B8 | 16 | 0.17 |
| B9 | 2 | 0.3 |
| B10 | 1 | 0.3 |
| B10 | 10 | 0.33 |
| B11 | 11 | 0.26 |
| B12 | 12 | 0.21 |
| B13 | 10 | 0.3 |
| B14 | 8 | 0.3 |

**A**

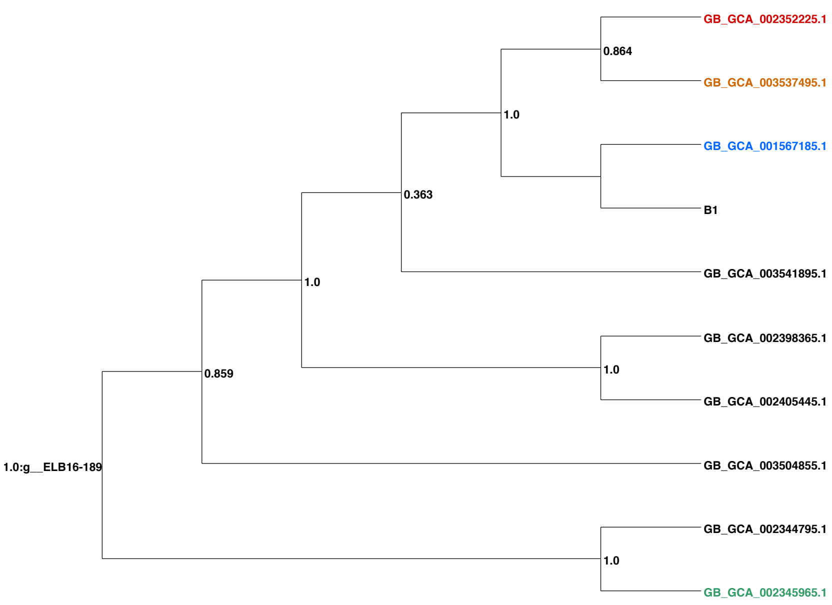


**B**

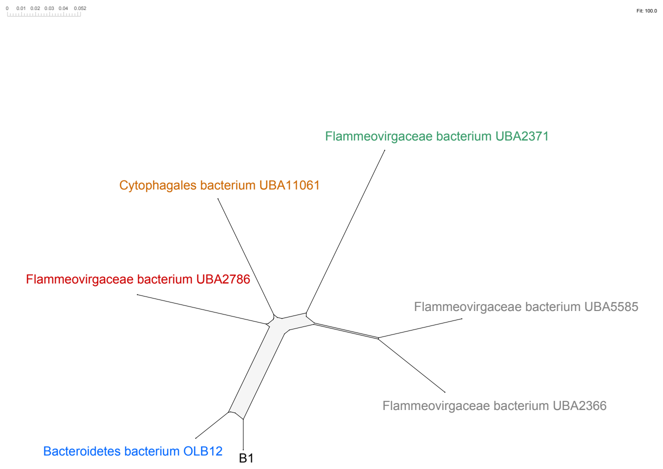


Figure S1: Phylogenetic tree computed by GTDB-Tk (A), and phylogenetic outline computed by SplitsTree5 (B) for the bin B1 - High quality draft genome of *Bacteroidetes bacterium* OLB12

**A**

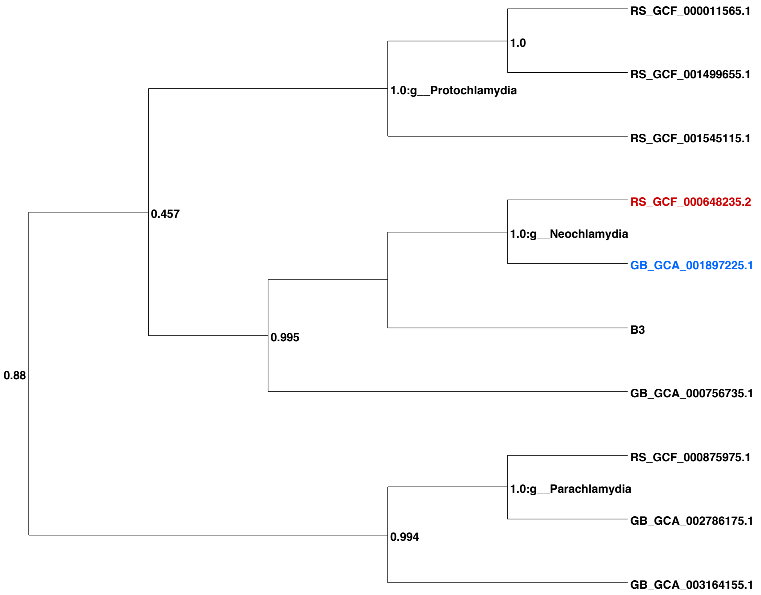


**B**

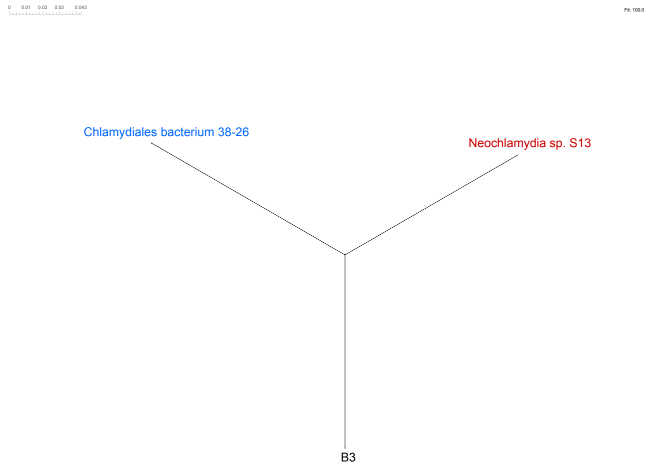


Figure S2: Phylogenetic tree computed by GTDB-Tk (A), and phylogenetic outline computed by SplitsTree5 (B) for the bin B3 - High quality draft genome of Chlamydiia

A

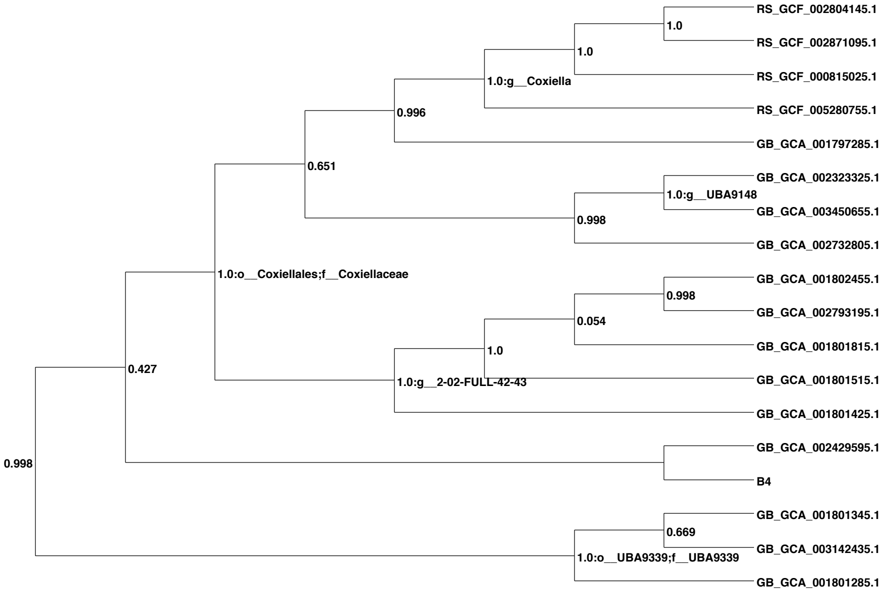


B

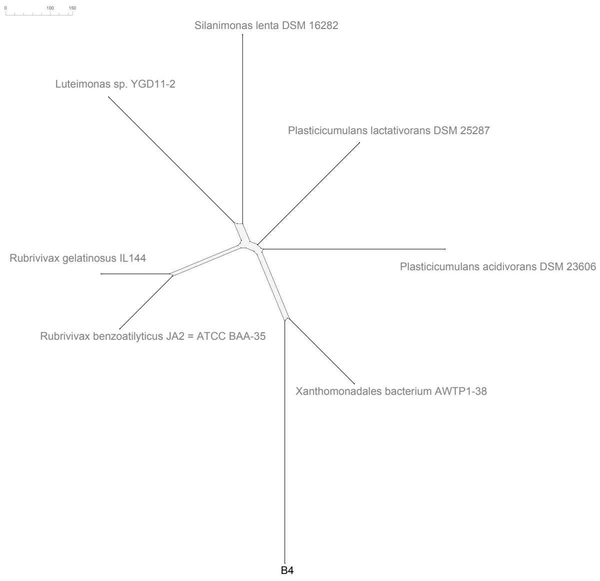


Figure S3: Phylogenetic tree computed by GTDB-Tk (A), and phylogenetic outline computed by SplitsTree5 (B) for the bin B4 - High quality draft genome of Gammaproteobacteria

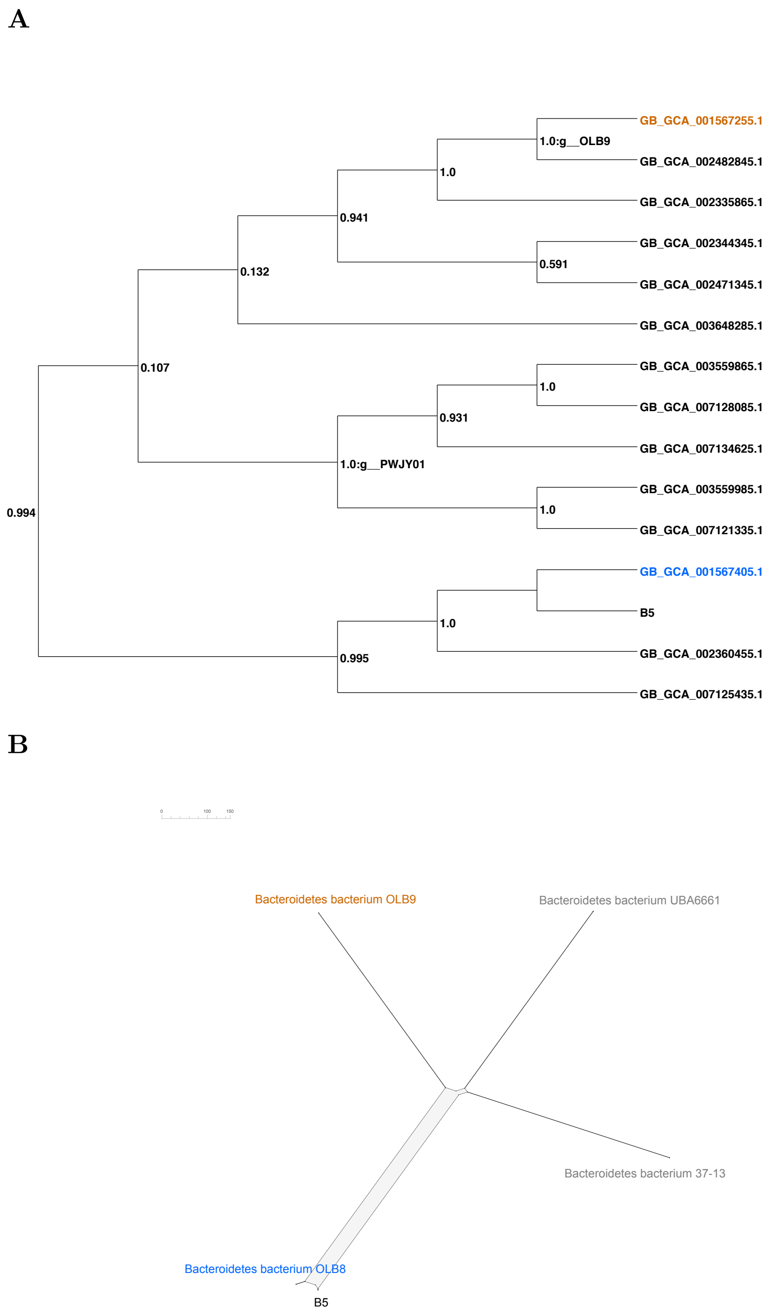


Figure S4: Phylogenetic tree computed by GTDB-Tk (A), and phylogenetic outline computed by SplitsTree5 (B) for the bin B5 - High quality draft genome of *Bacteroidetes bacterium* OLB8

**A**

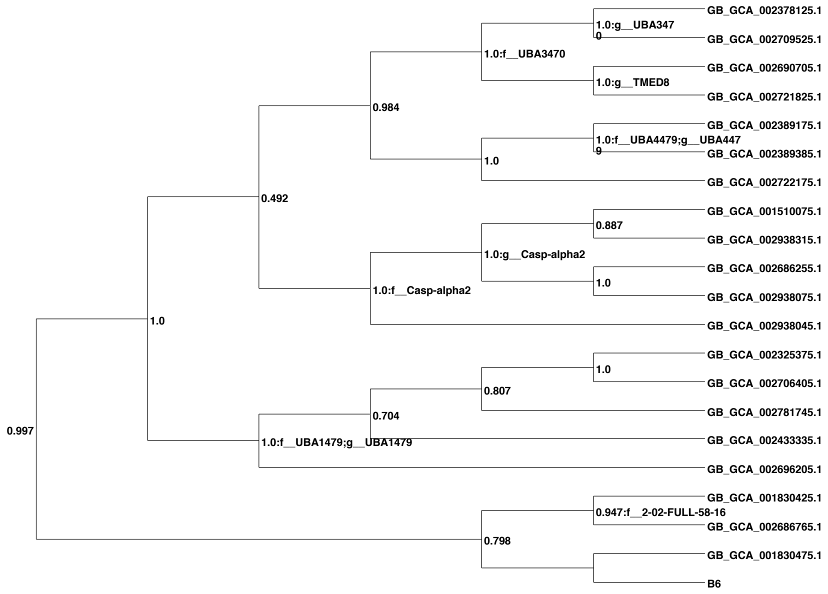


**B**

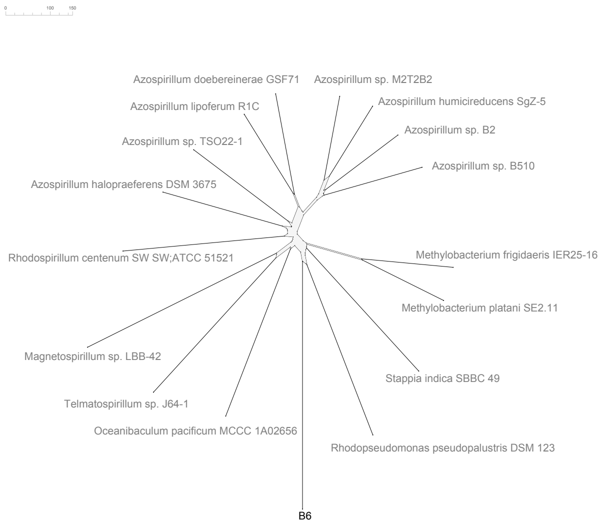


Figure S5: Phylogenetic tree computed by GTDB-Tk (A), and phylogenetic outline computed by SplitsTree5 (B) for the bin B6 - High quality draft genome of Rhodospirillales

**A**

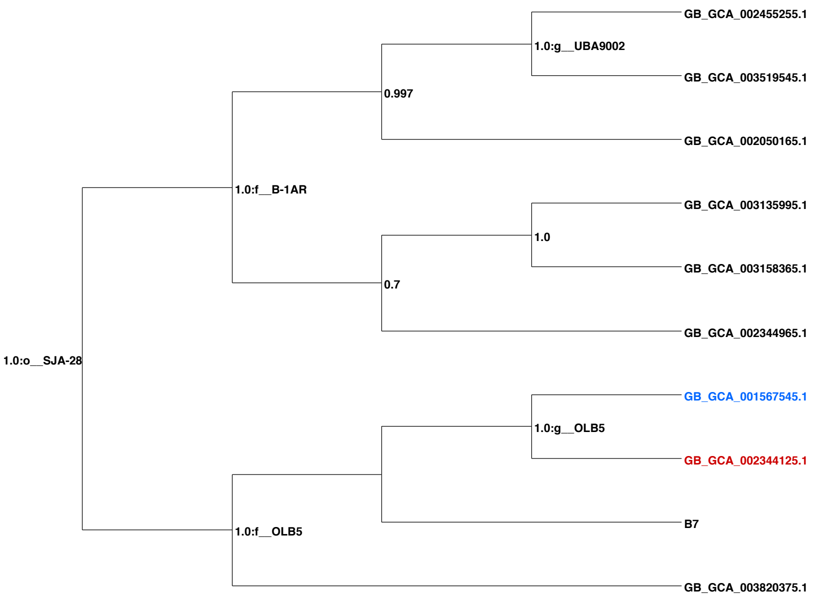


**B**

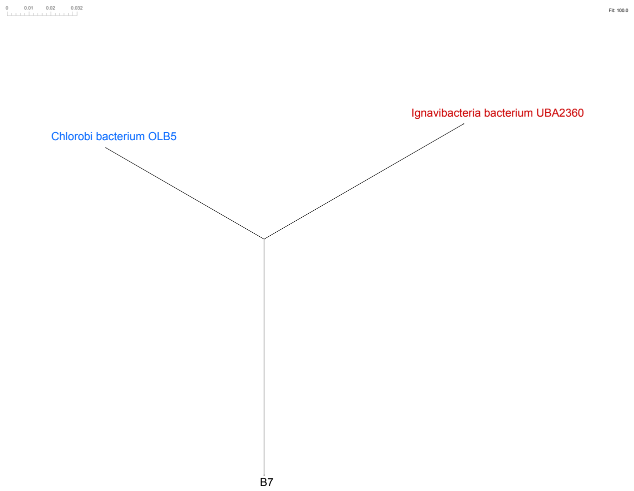


Figure S6: Phylogenetic tree computed by GTDB-Tk (A), and phylogenetic outline computed by SplitsTree5 (B) for the bin B7 - High quality draft genome of *Chlorobi bacterium* OLB5

**A**

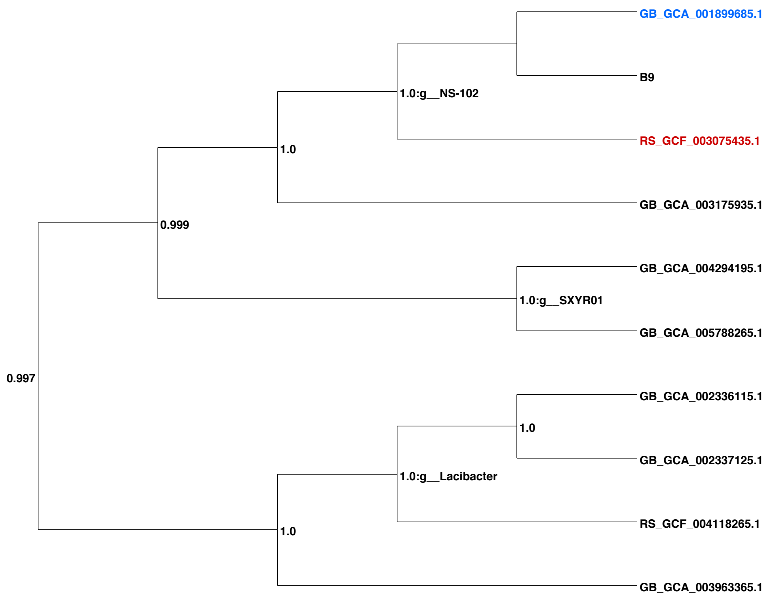


**B**

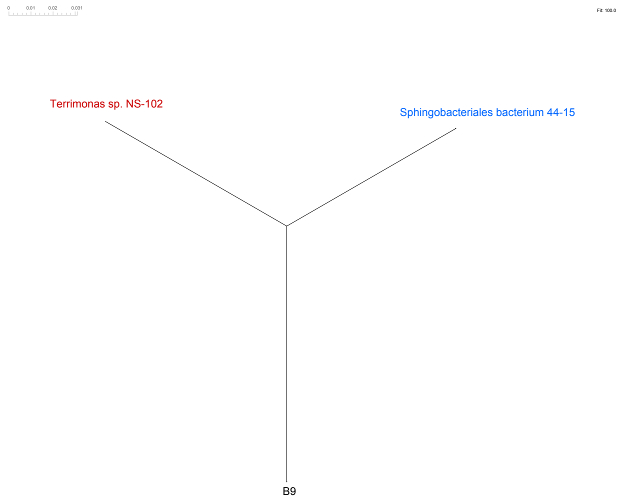


Figure S7: Phylogenetic tree computed by GTDB-Tk (A), and phylogenetic outline computed by SplitsTree5 (B) for the bin B9 - Medium quality draft genome of *Sphingobacteriales bacterium* 44-15

**A**

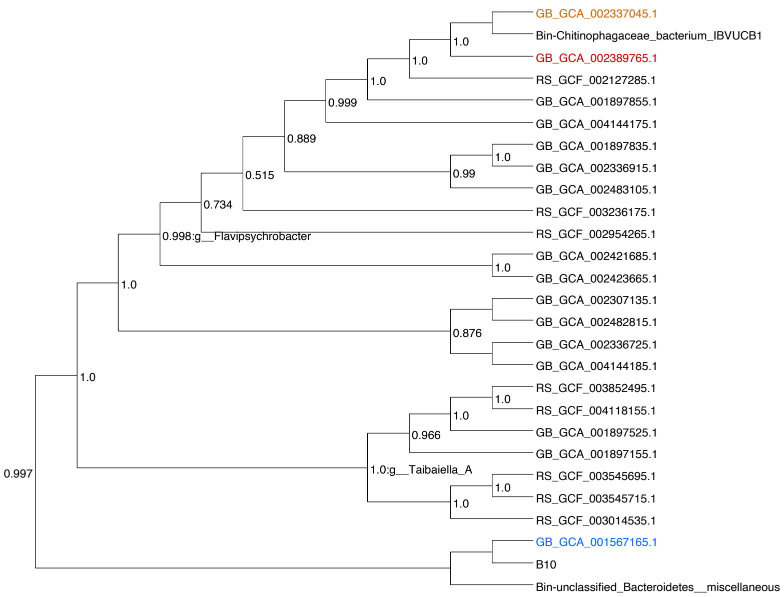


**B**

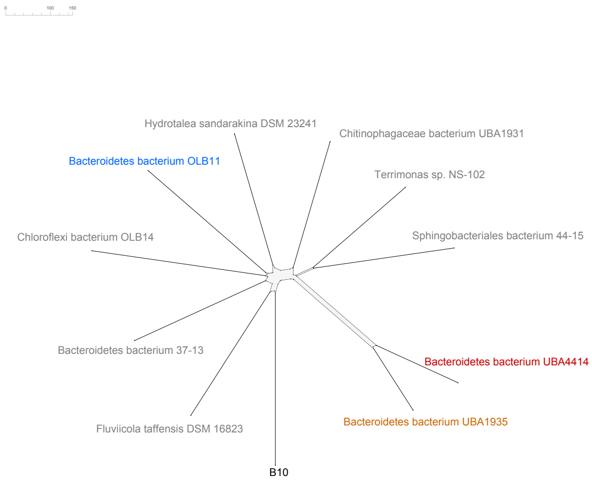


Figure S8: Phylogenetic tree computed by GTDB-Tk (A), and phylogenetic outline computed by SplitsTree5 (B) for the bin B10 - Medium quality draft genome of Bacteroidetes

**A**

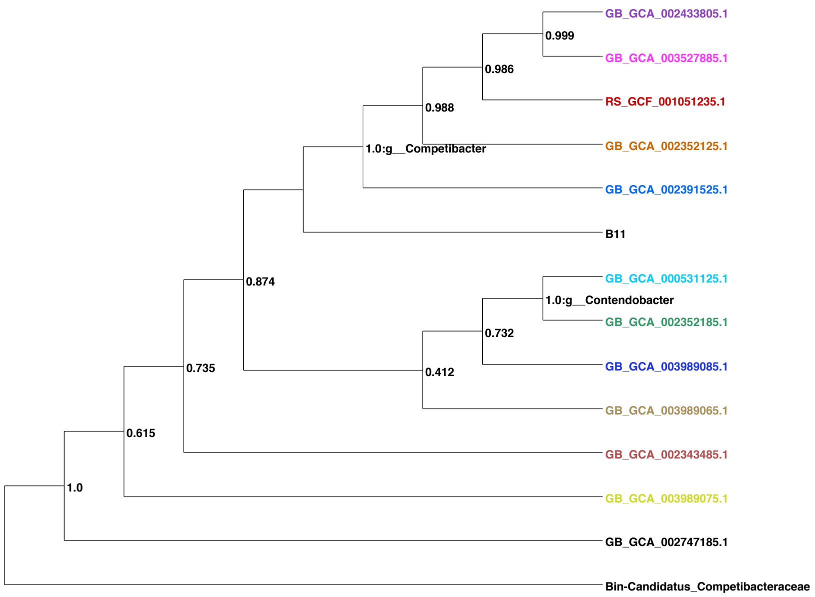


**B**

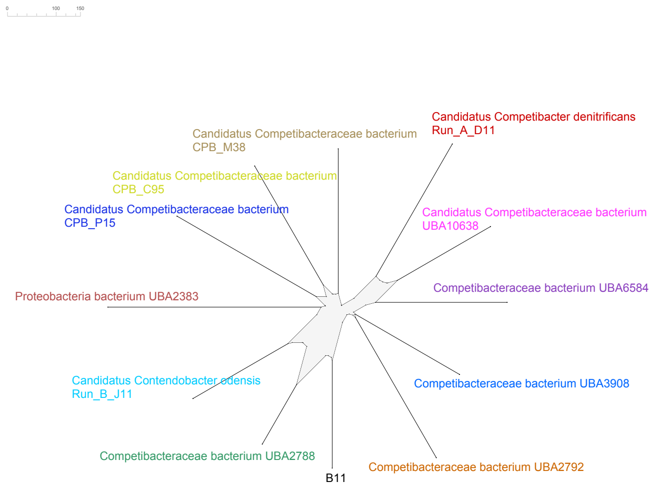


Figure S9: Phylogenetic tree computed by GTDB-Tk (A), and phylogenetic outline computed by SplitsTree5 (B) for the bin B11 - Medium quality draft genome of *Candidatus Contendobacter* B J11

**A**

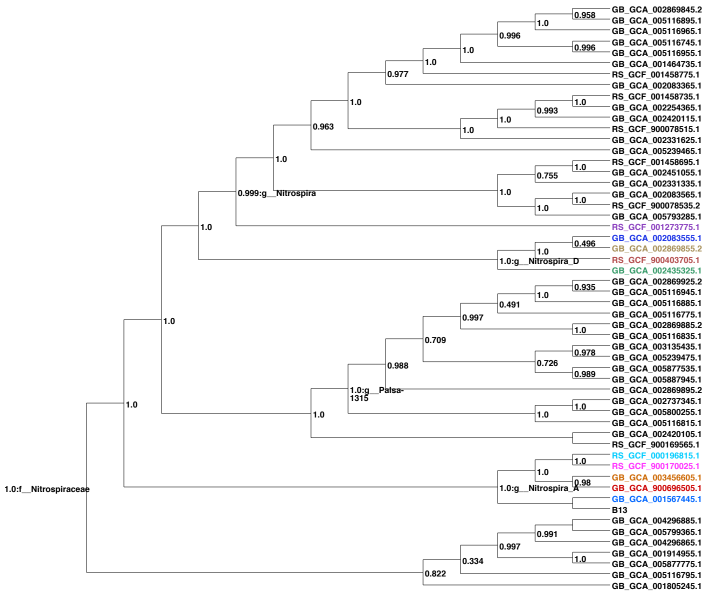


**B**

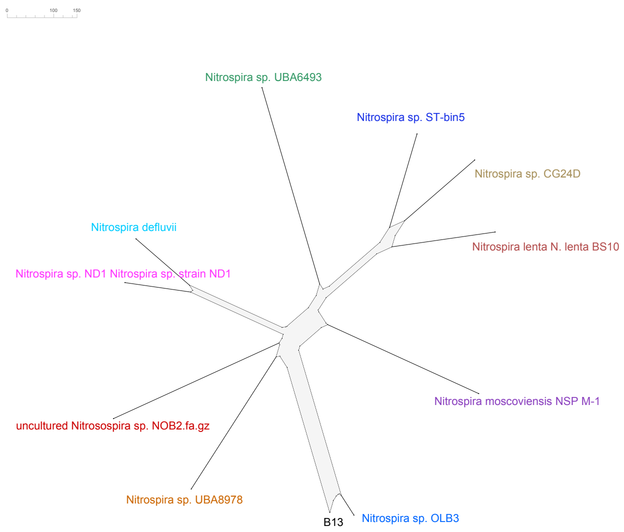


Figure S10: Phylogenetic tree computed by GTDB-Tk (A), and phylogenetic outline computed by SplitsTree5 (B) for the bin B13 - Low quality draft genome of *Nitrospira*

**A**

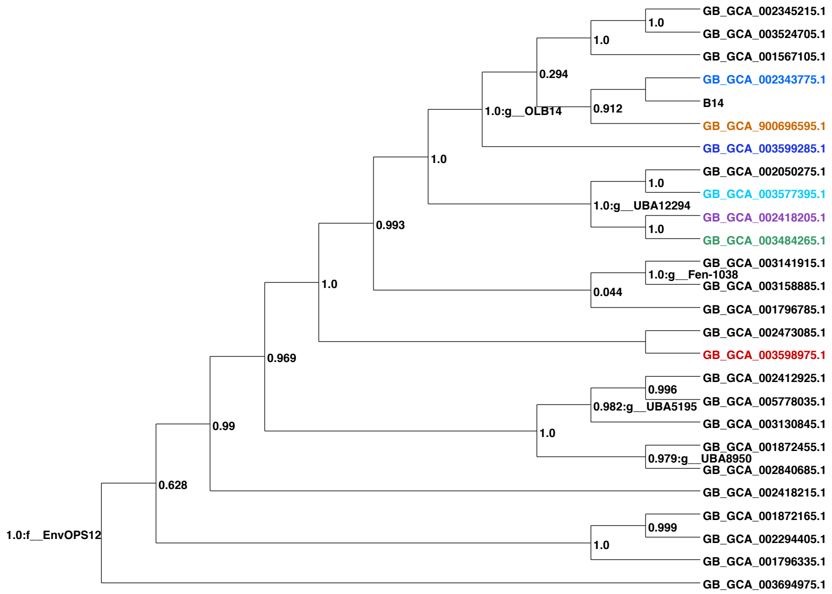


**B**

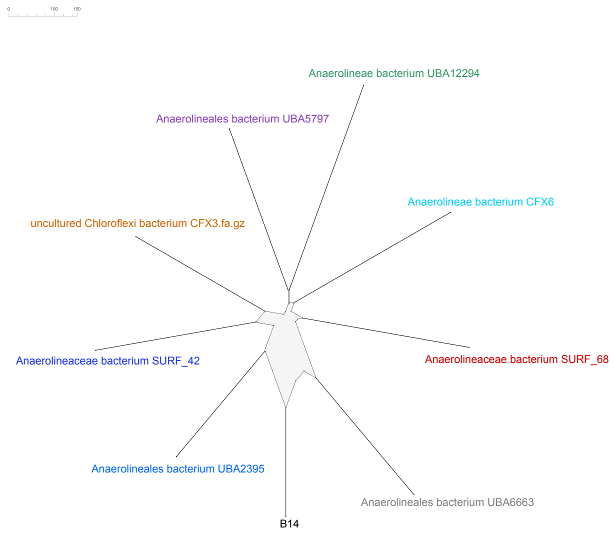


Figure S11: Phylogenetic tree computed by GTDB-Tk (A), and phylogenetic outline computed by SplitsTree5 (B) for the bin B14 - Low quality draft genome of Chloroflexi
